# Genome-wide analysis of PTR/POT transporters in *Candida* species and their functional characterization in the newly emerged pathogen *Candida auris*

**DOI:** 10.1101/2022.03.14.484144

**Authors:** Rosy Khatoon, Suman Sharma, Rajendra Prasad, Andrew M. Lynn, Amresh Prakash, Atanu Banerjee

## Abstract

The PTR or proton-dependent oligopeptide transporter (POT) family exploits the inwardly directed proton motive force to facilitate the cellular uptake of di- and tripeptides. Interestingly, representatives from the family have shown efficacy in delivering peptide-based antifungal derivatives in certain *Candida* species. Given the increased incidences of fungal infections by *Candida* species and the associated escalating orders of resistance against frontline antifungals, peptide derivatives are attractive therapeutic options. In that direction, the identification and characterization of PTR transporters serve as an essential first step in the translation of peptide-based antifungals as next-gen therapeutics. Herein, we present a genome-wide inventory of the PTR transporters in five prominent *Candida* species. Our study identifies 2 PTR transporters each in *C. albicans* and *C. dubliniensis*, 1 in *C. glabrata*, 4 in *C. parapsilosis*, and 3 in *C. auris*. Notably, despite all representatives retaining the conserved features seen in the PTR family, there exist two distinct classes of PTR transporters that differ in terms of their sequence identities and membrane topology. Further, we also evaluated the contribution of each PTR protein of the newly emerged multidrug-resistant *C. auris* in di-/tripeptide uptake. Notably, deletion of the PTR transporters encoded by BNJ08_003830 and BNJ08_005124 led to a marked reduction in the transport capabilities of several tested di-/tripeptides. Besides, BNJ08_005124 deletion also resulted in increased resistance towards the peptide-nucleoside drug Nikkomycin Z, pointing towards its predominant role in the uptake mechanism. Altogether, the study provides an important template for future structure-function investigations of PTR transporters in *Candida* species.

## INTRODUCTION

Pathogenic yeasts including *Candida albicans* and non-*albicans Candida* (NAC) species are successful human commensals and assume pathogenic status when exposed to compromised immunity (Sanches et al. 2019). The superficial infections caused by *C. albicans* and NAC species can also extend to disseminated bloodstream and deep-tissue infections (Sanches et al. 2019). While C. *albicans* and few NAC species (e.g., *C. glabrata, C. tropicalis, C. parapsilosis, C. dubliniensis, C. guilliermondii, C. krusei, and C. kefyr*) are the most common species affecting immunocompromised patients, more recently *C. auris* has established itself as a global health threat (https://www.cdc.gov/fungal/candida-auris/index.html) (Chowdhary et al. 2017; Taei et al. 2019). The biggest problem faced with this particular species is its ability to rapidly spread in hospital settings and the high order of resistance against the frontline antifungals (Lockhart et al. 2016; Chowdhary et al. 2017). An increase in the levels of antifungal resistance is also witnessed in *Candida albicans* and several other NAC species (Whaley et al. 2017). As a result, fungal research across the globe is currently directed towards the identification of alternative drug targets and therapeutics. Peptide transport serves as one of the primary routes for the acquisition of nitrogen and amino acids by cells, and peptide uptake systems are ubiquitously present in all organisms from bacteria to higher eukaryotes. (Newstead 2015). Two distinct classes of proton-coupled peptide transport systems are recognized in fungi: the peptide transporters (PTR/POT), which transport di- and tripeptides, and the oligopeptide transporters (OPT), which mediate the uptake of longer peptides (Dunkel et al. 2013; Becerra-Rodríguez et al. 2020). In other words, their specificity depends on the peptide length. Both these families belong to the larger Major Facilitator superfamily (MFS) of membrane-bound pumps (Garai et al. 2017). It was Paulsen and Skurray (1994) who first noted sequence similarities between different non-ABC-type, proton dependent transporters, like *Arabidopsis thaliana* nitrate transporter Chl1, mammalian PepT1, YhiP/dtpB transport system of *Escherichia coli*, oligopeptide transporter DtpT from *Lactococcus lactis*, and the *S. cerevisiae* peptide transporter Ptr2 and hence designated the group as Proton-dependent Oligopeptide Transport (POT) family (Paulsen and Skurray 1994). The authors also predicted a shared topological domain consisting of 12 putative a-helical membrane-spanning segments, with two unique conserved motifs not known to be present in any other class of proteins at that time. Later, some research groups did not consider the POT family designation completely correct for the set of proteins and proposed the Peptide Transport family (PTR) to refer to a group of non-ABC type, proton-dependent oligopeptide transporters (Steiner et al. 1995). However, currently, the PTR or POT designations are interchangeably used in literature and refer to the proton-dependent oligopeptide transporter family with di- and tripeptides as the substrates (Newstead 2015). Interestingly, this group of transporters has been identified in both eukaryotes and prokaryotes, except archaea, which can be partly ascribed to the fact that peptides are usually not utilized as nutrient sources in extreme environments in which the archaeal members thrive (Newstead 2017). In *S. cerevisiae*, the PTR transport system comprises of a single transporter ScPtr2 which is the best-known member of the PTR family with homologs reported in *Candida albicans* (Perry et al. 1994). An analysis of peptide specificity of ScPtr2 revealed its higher affinity for di- and tripeptides with aromatic, branched, or basic amino acids and a relatively lower affinity for negatively charged amino acids, glycine, and proline (Ito et al. 2013). In *C. albicans* two dipeptide/tripeptide transporters of the PTR family have been identified (Dunkel et al. 2013). One encoded by orf19.2583 is designated as CaPtr2 (Dunkel et al. 2013). The second protein encoded by orf19.6937 is more similar to ScPtr2 in terms of its primary structure and is designated as CaPtr22 (Dunkel et al. 2013). It was also noticed by Dunkel and colleagues that CaPtr22 has a broader substrate spectrum with an increased diversity of tripeptides that can be transported by it (Dunkel et al. 2013). Their counterparts in other *Candida* species have not been identified and/or investigated in detail and form the basis of this study. The primary reason why this class of protein requires investigation is that this class of proton-coupled transporters is also known for its pharmacological importance since they aid in the uptake of drugs that have steric resemblance with substrate peptides (Newstead 2015). Interestingly, in *C. albicans*, the di-/tripeptide transporters belonging to the PTR/POT family have demonstrated the ability to import antifungal peptides and thus open avenues for exploiting these membrane-bound pumps as antifungal delivery systems (Schielmann et al. 2017; Skwarecki et al. 2018). Furthermore, the PTR transporters in humans, PepT1 and PepT2, besides catalyzing the uptake of dietary peptides, also recognize and transport many drug molecules with steric resemblance towards peptides like β-lactam antibiotics cefadroxil and cefalexin (Terada et al. 1997). Since peptide-based therapeutics are known to be relatively refractory to resistance development, these systems can be constituent of novel treatment regimens. However, a deeper understanding of the structure and function of these proteins is required to fully utilize their potential. With this aim, the current study is aimed at the identification of PTR/POT transporters in predominant *Candida* species to pave the way for exploiting these proteins as efficient antifungal delivery systems. We also attempted to functionally characterize these systems in the newly emerged *C. auris* to encourage further structure-function studies.

## MATERIALS AND METHODS

### BLAST analysis

To identify the PTR transporters in *Candida* species, BLAST+ feature within the Candida Genome Database (CGD) (Inglis et al. 2012) was utilized using the ScPtr2 protein sequence as the query and *C. albicans, C. dubliniensis, C. glabrata, C. parapsilosis*, and *C. auris* as the target genomes. BLAST was run using the default settings and results were analyzed based on the E-value and normalized alignment scores (bits).

### Identification of Protein domains

Protein sequences of all the predicted Ptr candidates were retrieved from CGD and a combined FASTA file was submitted as an input to the DomainViz webserver (Schläpfer et al. 2021). Default parameters were utilized for the job.

### Multiple sequence alignment and clustering sequence identity matrix construction

Multiple sequence alignment (MSA) of the identified protein sequences was constructed using PSI/TM-Coffee (http://tcoffee.crg.cat/apps/tcoffee/do:tmcoffee). Within the homology search options, sequence type was set to “transmembrane” and UniRef100 was selected for homology extension. The alignment output was downloaded in the FASTA format and visualized in the Jalview alignment editor (Waterhouse et al. 2009) for the identification of conserved motifs. The same MSA was also used to generate the percent identity matrix using BioEdit sequence alignment editor. The resultant sequence identities were used to generate the clustering sequence identity matrix with the help of Seaborn data visualization library (https://seaborn.pydata.org/generated/seaborn.clustermap.html)in Python.

### Phylogenetic tree construction

The MSA generated above was utilized to generate a phylogenetic tree in MEGA-X (Kumar et al. 2018) using the Maximum likelihood method. The bootstrap consensus tree inferred from 1000 replicates was taken to represent the evolutionary history of the taxa analyzed. The generated tree data was exported in Newick format and submitted to iTOL (https://itol.embl.de/) (Letunic and Bork 2021) to generate the final tree figure.

### Membrane topology analysis

To analyze the membrane topology, the FASTA file containing protein sequences was submitted to TOPCONS (https://topcons.cbr.su.se) (Tsirigos et al. 2015). The sequence file along with the membrane topology data from TOPCONS was then submitted to Protter (https://wlab.ethz.ch/protter/start/) (Omasits et al. 2014) to generate the membrane topology diagrams for the identified proteins.

### Materials

All routine chemicals used in the study were purchased from SRL Pvt. Ltd., Mumbai, India. The dipeptide, tripeptides, and Nikkomycin Z were procured from Sigma-Aldrich Co. (St. Louis, MO). Nourseothricin Sulfate was purchased from Gold biotechnology Inc. (USA). Oligonucleotides were purchased from Bioserve Biotechnologies India Pvt. Limited.

### Strains and culture conditions

The *C. auris* strain CBS10913T was obtained from the CBS-KNAW fungal culture collection of the Westerdijk Fungal Biodiversity Institute, Utrecht, Netherlands, and reported in our previous study (Wasi et al. 2019). The yeast strains were maintained and cultured in the YEPD medium (yeast extract, peptone, and dextrose) from HiMedia Laboratories, Mumbai, India. For the peptide uptake assays, Yeast Carbon base (YCB) media (HiMedia Laboratories, Mumbai, India) was used along with the dipeptides or tripeptides as the nitrogen source. All the yeast strains used in the study are listed in **Supplementary table 1**.

### Construction of *PTR* deletion mutants in CBS10913T

For the construction of deletion mutant, we used the fusion PCR-based deletion strategy. Firstly, 5’-and 3’-UTR regions (nearly 600-700 bps) of the genes (BNJ08_004188, BNJ08_003830, and BNJ08_005124) were amplified from genomic DNA of *C. auris* CBS10913T. The selection marker *NAT1* gene with FRT sites was also PCR amplified in two halves from plasmid pRK625. Both 5’-and 3’-UTR amplified products were fused to one-half each of the NAT1 gene fragments to generate the fusion product. The fused PCR products were then co-transformed into the CBS10913T (WT) strain using electroporation strategy and positive transformants were selected on YEPD agar medium containing 200 μg/ml nourseothricin. The gene deletions were confirmed by gene-specific PCRs from the genomic DNA of transformants. The same was also finally confirmed through DNA sequencing. All the oligonucleotides used for gene deletions and their confirmations are enlisted in **Supplementary table 2**.

### Peptide uptake assay

For the peptide uptake assay, first overnight primary cultures of WT and mutants grown in YEPD media were centrifuged and washed with sterile MQ. Next, for each strain, OD_600_ = 0.2 was set in 1 ml of Yeast carbon base (YCB) containing 1 mg of each di-/tripeptide. The culture tubes were then incubated at 30°C for 48 hours with shaking. Cell turbidity was determined at 600 nm wavelength using a spectrophotometer.

### Nikkomycin Z sensitivity assay

Nikkomycin Z sensitivity in liquid medium was determined in 96-well microplate format. Herein, two-fold serial dilution of the compound was first prepared in YNB (Yeast nitrogen base) medium containing amino acids (YNB+AA). This medium comprised of 0.67 % YNB (Yeast nitrogen base) without amino acids (Difco, Becton, Dickinson and Company, MD, USA), 0.2% dropout mix without uracil (Sigma-Aldrich Co.), 0.002% uracil (SRL), and 2% glucose (HiMedia). The medium was then incubated with logarithmic phase cells (∼10^4^ cells/ml) at 30°C for two days. Cell turbidity was determined at 595 nm wavelength using a microplate reader (BioRad).

For serial dilution spot assay, yeast suspensions were prepared in 0.9% NaCl solution and serially diluted five-fold. A 3 µl aliquot from each dilution was spotted onto YNB+AA agar plates with or without Nikkomycin Z (2µg/ml) and incubated for 48 h at 30°C.

## RESULTS

### Identification of PTR/POT transporters in *Candida* species

To identify the PTR transporters in *Candida* species, we utilized the Candida Genome Database (CGD) Multi-genome search using BLAST+ feature, wherein, we supplied the ScPtr2 sequence as the query and *C. albicans, C. dubliniensis, C. glabrata, C. parapsilosis*, and *C. auris* as the target genomes. The analysis revealed 2 PTR transporters each in *C. albicans* and *C. dubliniensis*, 1 in *C. glabrata*, 4 in *C. parapsilosis*, and 3 in *C. auris*. **Table 1** provides the details of the proteins and the strain IDs, and **Supplementary file 1** summarizes the BLAST results. The *C. glabrata* PTR transporter CAGL0G02453g showed the maximum percent identity to the query (∼71%) followed by CPAR2_503540 of *C. parapsilosis* and Cd36_83780 of *C. dubliniensis*, both having around 59% identity. The other 3 candidate PTR proteins of *C. parapsilosis* (CPAR2_800760, CPAR2_804430, and CPAR2_804220) showed relatively lower identities (in the range of 31-34%) to ScPtr2. The second *C. dubliniensis* candidate, Cd36_26850 also showed a relatively lower identity of ∼ 28% to ScPtr2. For *C. albicans*, we observed that CaPtr22 is much more identical to ScPtr2 (∼58% identity) than CaPtr2 (∼28% identity), as reported earlier too (Dunkel et al. 2013). Interestingly, all the three PTR candidate proteins in *C. auris* showed around 58-62% identity to ScPtr2. From these results, one point emerged that in certain *Candida* species, namely *C. albicans, C. parapsilosis, C. dubliniensis*, there exists one PTR transporter which is very similar to ScPtr2, while the other candidate protein(s) show low sequence identity to ScPtr2.

**Table 1:**
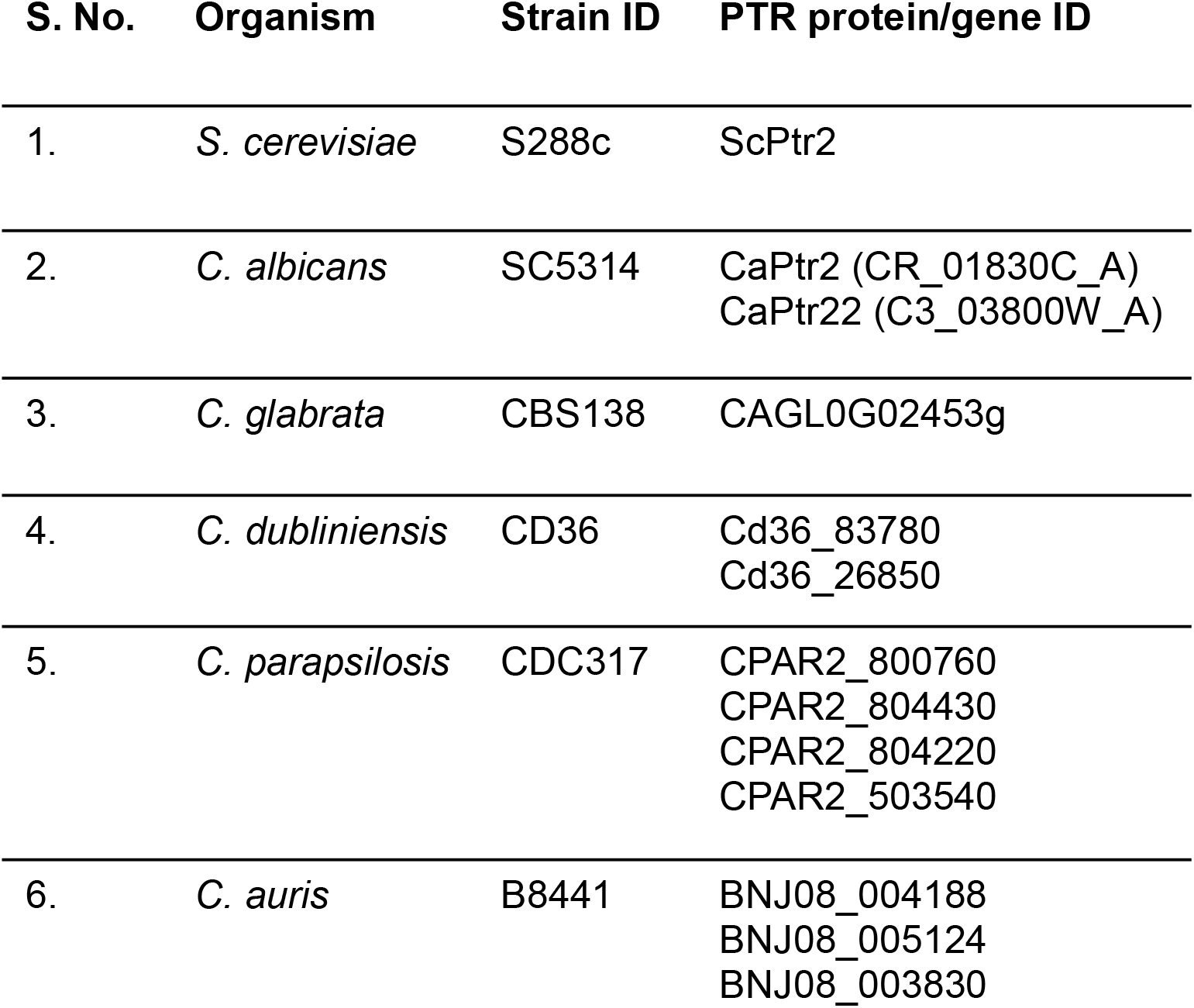
Putative PTR transporter encoding genes identified in the genome of *Candida* species.

| S. No. | Organism | Strain ID | PTR protein/gene ID |
| --- | --- | --- | --- |
| 1. | *S. cerevisiae* | S288c | ScPtr2 |
| 2. | *C. albicans* | SC5314 | CaPtr2 (CR_01830C_A)<br>CaPtr22 (C3_03800W_A) |
| 3. | *C. glabrata* | CBS138 | CAGL0G02453g |
| 4. | *C. dubliniensis* | CD36 | Cd36_83780<br>Cd36_26850 |
| 5. | *C. parapsilosis* | CDC317 | CPAR2_800760<br>CPAR2_804430<br>CPAR2_804220<br>CPAR2_503540 |
| 6. | *C. auris* | B8441 | BNJ08_004188<br>BNJ08_005124<br>BNJ08_003830 |

To further confirm the status of the identified proteins as PTR/POT family transporters, we exploited the DomainViz web server that uses PFAM (Finn et al. 2016) and Prosite databases (Sigrist et al. 2013) for domain searching and displays the extent of domain conservation along two dimensions: positionality and frequency of occurrence in the input protein sequences (Schläpfer et al. 2021). As evident from **Supplementary figure 1**, PTR2 (PF00854) is the most represented PFAM domain, with all the submitted sequences containing it. One protein also shows MFS_1 PFAM domain, however, HMMSCAN based search of the query (CAGL0G02453g) against PFAM domains results in its exclusion by the PFAM post-processing. Further, all submitted sequences contain either PTR2_1 or PTR2_2 PROSITE signatures (**Supplementary figure 1)**. Overall, it could be concluded that all the selected proteins from our BLAST analysis belong to the proton-dependent oligopeptide transporters (POT/PTR family).

### Multiple sequence alignment (MSA) to reveal the presence of conserved motifs characteristic of the PTR/POT family

To gain further sequence-related insights, we performed multiple sequence alignment of the identified proteins using PSI/TM-Coffee (http://tcoffee.crg.cat/apps/tcoffee/do:tmcoffee) (Floden et al. 2016) which allows efficient alignment of transmembrane proteins using homology extension on reduced databases. The tool outperforms many other available tools available for the purpose (Floden et al. 2016). We also verified the constructed MSA using another newly developed tool for membrane protein alignment, TM-Aligner (Bhat et al. 2017). Both methods produced comparable results. We visualized the generated MSA in Jalview alignment viewer (Waterhouse et al. 2009) and performed further analysis within this application only. One interesting observation made with the MSA was that two of the *C. parapsilosis* transporters, CPAR2_804430 and CPAR2_804220 are 99.8% identical with a difference of only a single amino acid, whereby isoleucine in the former is replaced by threonine at 223 position (data not shown).

Contrary, to other membrane transporter families, the PTR family representatives show high order of sequence similarity at the level of transmembrane helices (TMHs) (Steiner et al. 1995; Newstead 2015). The constructed MSA also revealed many conserved blocks of amino acids (data not shown). A total of three typical sequence motifs have been reported in the PTR family. One motif is referred to as the ExxERFxYY motif on TMH1 (Newstead 2015). The other two relatively less well-conserved motifs are denoted as the PTR2-1 and PTR2-2 (Newstead 2015). Mutational studies have also revealed their relevance in the transport mechanism of PTR/POT transporters (Hauser et al. 2005; Solcan et al. 2012; Newstead 2015). **Figure 1** shows a color-coded MSA (based on identity) of the PTR candidates in *Candida* species as well as *S. cerevisiae* highlighting the regions of the three conserved motifs. It also depicts the conservation scores for each alignment column and consensus motifs. As evident from the sequence logo, all three motifs are well conserved across all the representatives.

**Figure 1:**
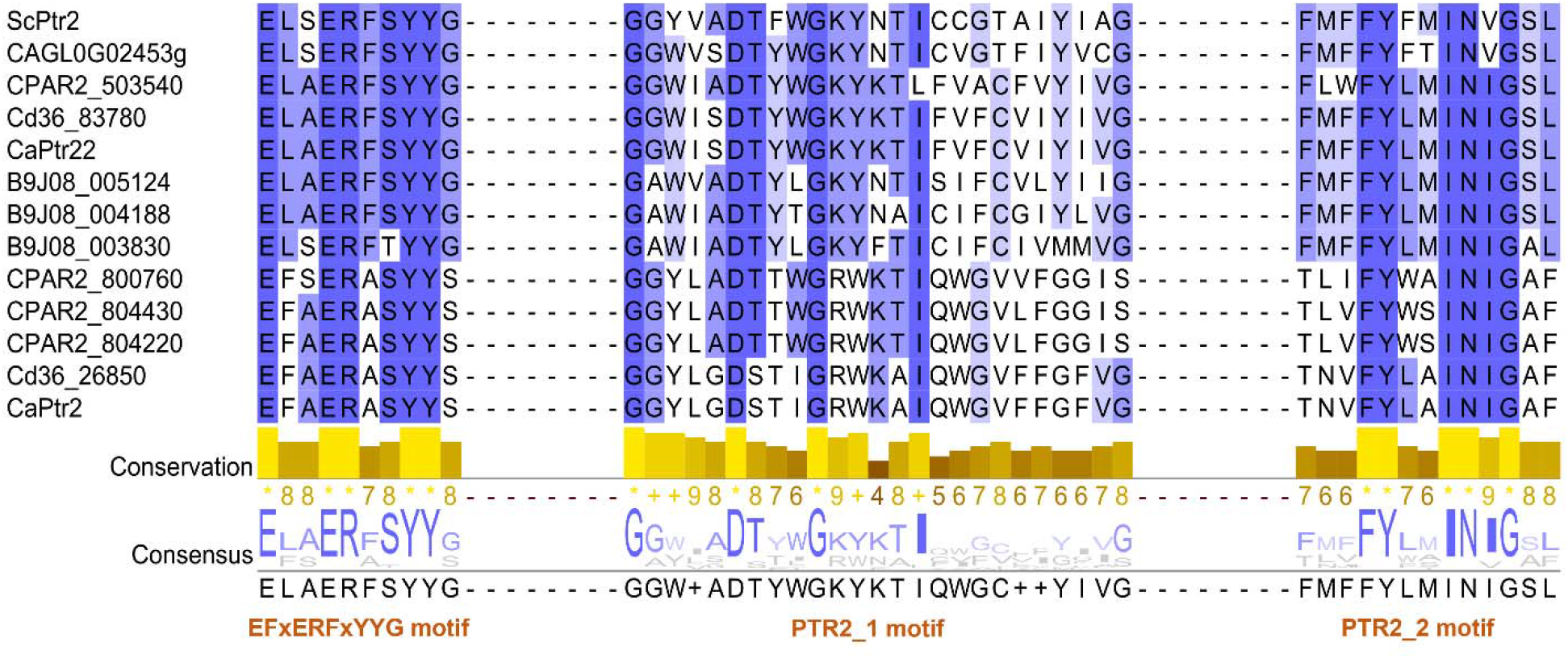
Snapshot of Multiple sequence alignment of all the putative Ptr transporter protein sequences constructed using PSI/TM-Coffee and visualized in Jalview highlighting the conservation of the three known motifs present in the PTR family.

### Construction of a clustering identity matrix to identify the sequence relatedness between different members

As described previously, several *Candida* species contained PTR protein(s) which show poor sequence identity to ScPtr2. To better understand this heterogeneity amongst the different representatives, we constructed a clustering sequence identity matrix based on the MSA generated above. As shown in **Figure 2A**, two major clusters of proteins emerge from the data. While one comprises of ScPtr2, CaPtr22, all three PTR transporters of *C. auris* (BNJ08_004188, BNJ08_005124, and BNJ08_003830). CAGL0G02453g transporter of *C. glabrata*, Cd36_83780 of *C. dubliniensis* and CPAR2_503540 of *C. parapsilosis*. The second cluster comprised of the remaining proteins: CaPtr2 of *C. albicans*; CPAR2_800760, CPAR2_804430, and CPAR2_804220 of *C. parapsilosis*; and Cd36_26850 of *C. dubliniensis*. This observation suggests that the low sequence identity that exists between the *C. albicans* PTR transporters (∼26%) extends to many other *Candida* species like *C. parapsilosis* and *C. dubliniensis*, wherein some protein(s) share low similarity amongst themselves. Overall, it could be said that within the PTR family of *Candida* species, two distinct classes of proteins exist, which will be referred to from here onwards as ScPtr2-like and ScPtr2-unlike groups. As mentioned before, the functional analysis of CaPtr2 and CaPtr22 also suggested a difference in the substrate repertoire of these two proteins (Dunkel et al. 2013). It is plausible that such a difference in substrate specificity also occurs among the PTR transporters in other *Candida* species. The question however is the need of having two distinct classes of PTR proteins.

**Figure 2:**
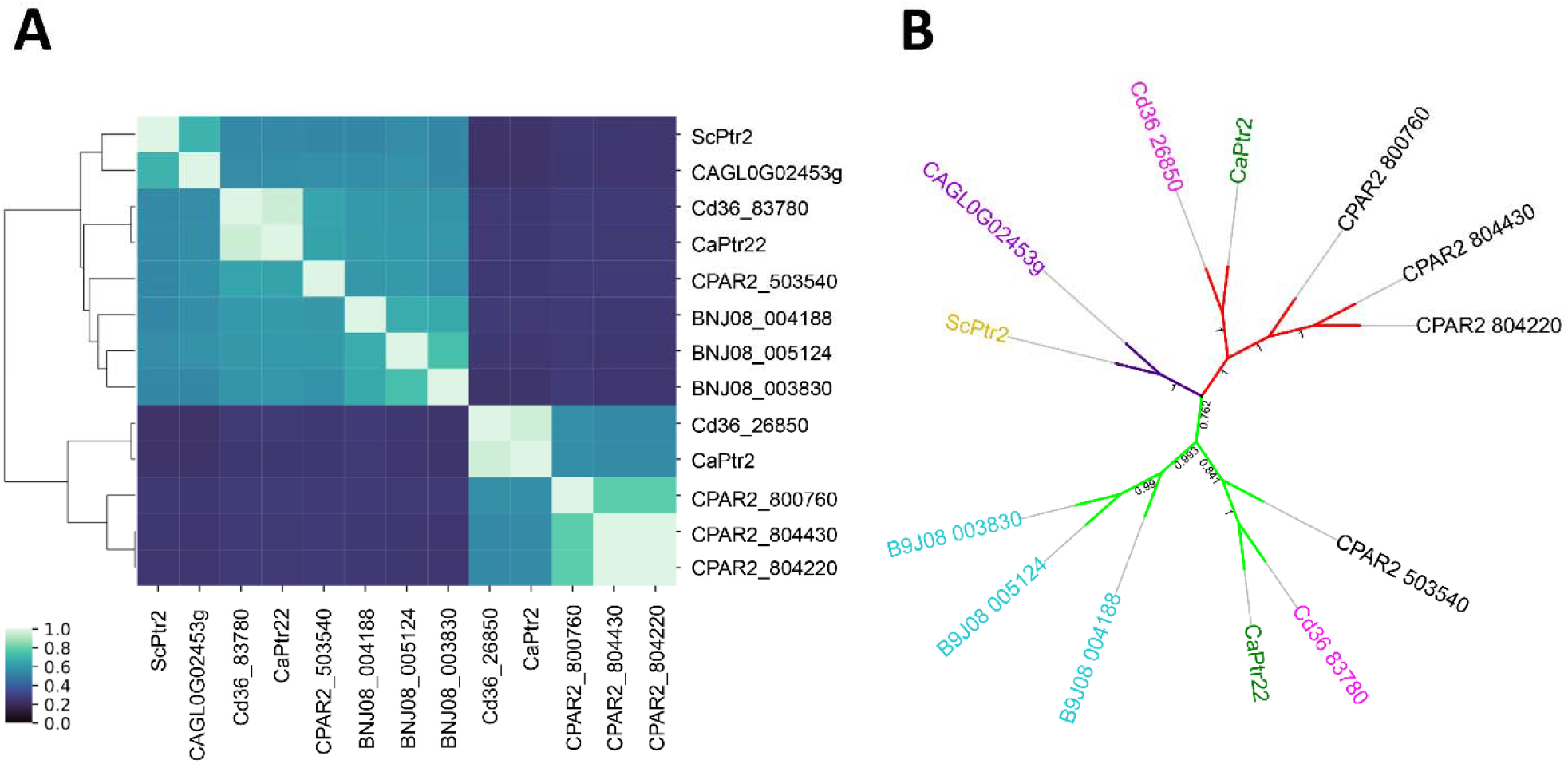
**(A)** Clustering sequence identity matrix of the Ptr protein sequences prepared using Seaborn library in Python. The input identity values for the plot were calculated from MSA generated using BioEdit tool. **(B)** Phylogenetic tree of the *Candida* PTR transporters constructed using the Maximum Likelihood method and Jones et al. w/freq. model. The bootstrap consensus tree inferred from 1000 replicates is taken to represent the evolutionary history of the taxa analyzed. The percentage of replicate trees in which the associated taxa clustered together in the bootstrap test (1000 replicates) are shown next to the branches. Initial tree(s) for the heuristic search were obtained automatically by applying Neighbor-Join and BioNJ algorithms to a matrix of pairwise distances estimated using the JTT model, and then selecting the topology with superior log likelihood value. Evolutionary analyses were conducted in MEGA X and further modified and visualized in iTOL.

### Phylogenetic analysis of the PTR representatives from *Candida* species

To obtain an evolutionary perspective of the results obtained so far, we performed phylogenetic analysis of the PTR candidates using the Maximum likelihood method Bootstrap values were calculated using 1,000 reiterations. The topology of the phylogenetic tree showed that the PTR transporters fall into three major groups **(Figure 2B)**. ScPtr2 and CAGL0G02453g are clustered together and form group 1. Group 2 comprises of CaPtr22, Cd36_83780, CPAR2_503540, BNJ08_004188, BNJ08_005124 and BNJ08_003830. Group 3 comprises the same set of proteins revealed in the sequence identity matrix to be showing remote sequence identity to ScPtr2, viz., CaPtr2, CPAR2_800760, CPAR2_804430, CPAR2_804220, and Cd36_26850. Overall, the grouping resembled the clustering obtained in the hierarchical sequence identity matrix. The obtained topology supports the results obtained from the abovementioned sequence analysis that suggested a distinct class of transporters within the PTR family which shows poor homology to ScPtr2.

### Membrane topology analysis of the PTR transporters

Given the results obtained before, we next chose to analyze the topology of the transporters to see if there is any distinction in the way the membrane, intracellular and extracellular segments are arranged within the PTR transporters. We utilized the TOPCONS tool for membrane topology prediction which is a widely used program for consensus prediction of membrane protein topology. TOPCONS combines an arbitrary number of topology predictions into one consensus prediction (Tsirigos et al. 2015) and thus plausibly identifies the number of TMHs and their arrangement accurately. As shown in **Figure 3**, all the proteins contain 12 TMHs with both the N- and C-terminals towards the intracellular region. The membrane topology also suggested that ScPtr2-like group has some notable differences when compared to the ScPtr2-unlike group as shown in **Figure 3A** and **3B**, respectively. The length of the central cytoplasmic loop (CCL) and Extracellular loop 1 (ECL1) is much longer in the ScPtr2-unlike group compared to the ScPtr2-like group. While the length of CCL in the ScPtr2-like group members is around 53 amino acids, it is around 70 amino acids in the ScPtr2-unlike group members (**Figure 4A**). Similarly, the average length of ECL1 is 28 and 42 amino acids in the former and latter groups, respectively (**Figure 4B**). Therefore, the membrane topology in the two groups also appears to be distinct, albeit in a subtle manner.

**Figure 3:**
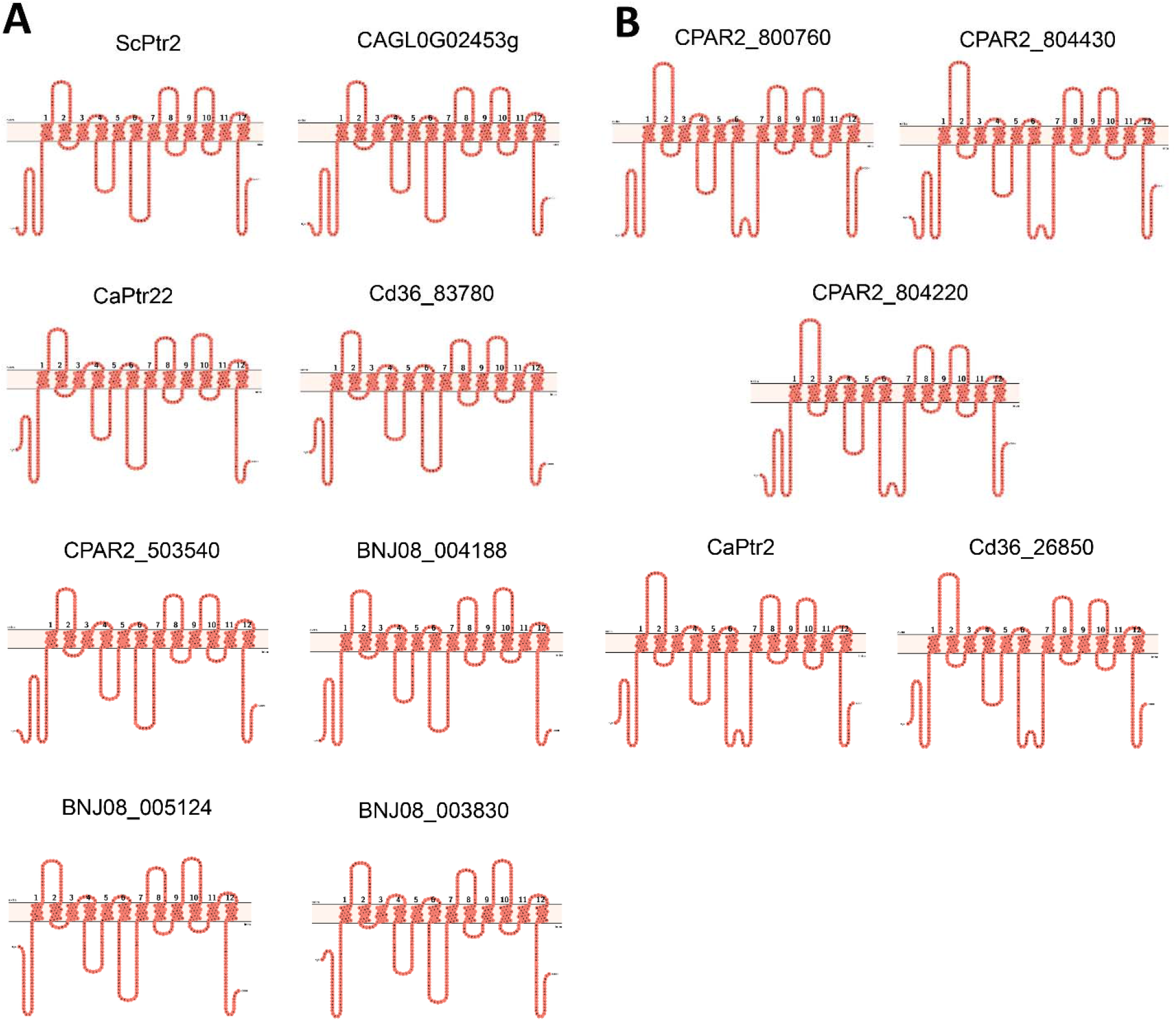
Membrane topology diagrams of all Ptr transporters prepared using Protter. The TM, extracellular and intracellular regions were predicted using TOPCONS and supplied as input to Protter. ScPtr2-like and ScPtr2-unlike groups are shown in panels **(A)** and **(B)**, respectively.

**Figure 4:**
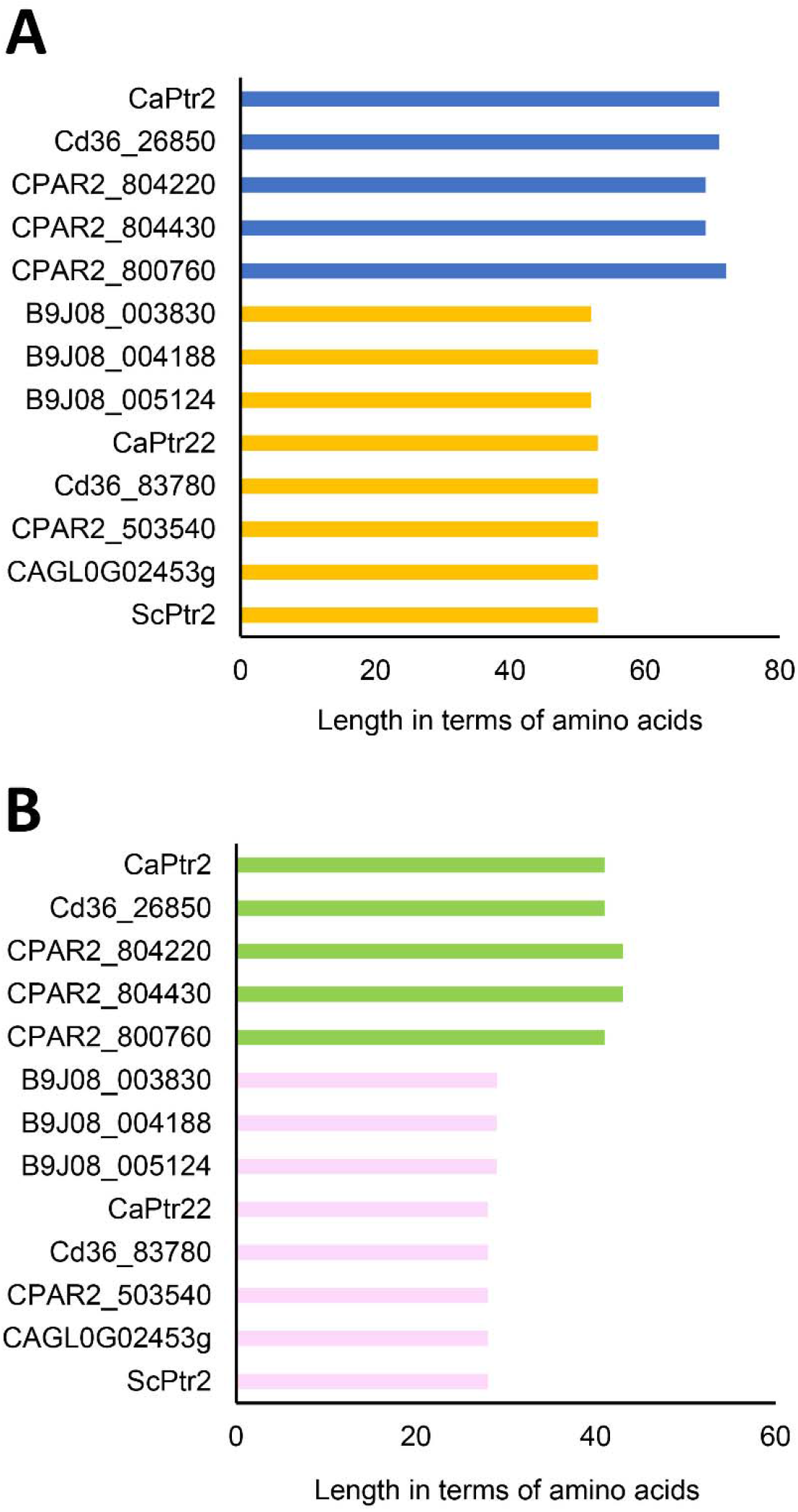
Bar-graphs representing the length of **(A)** Central cytoplasmic loop (CCL1) **(B)** Extracellular loop 1 (ECL1) in the Ptr proteins predicted using TOPCONS.

### Functional characterization of PTR transporters of *Candida auris*

The recent emergence of the multidrug-resistant pathogen *Candida auris* has triggered the search for alternative therapeutics. As explained before, PTR systems with their ability to import dipeptide antifungal conjugates are poised to serve as delivery systems for antifungals. Along similar lines, peptide-based antifungal compounds have recently shown promising antifungal activities (Bentz et al. 2021; Shahi et al. 2022). Given that PTR transporters are involved in their uptake (Yadan et al. 1984; Schielmann et al. 2017) and such proteins show increased expression in many clinical isolates (Muñoz et al. 2018; Shahi et al. 2022), we chose to characterize all PTR transporters encoded in the genome of *C. auris*. We constructed deletion mutants for all three identified PTR transporter encoding genes, viz. (BNJ08_004188, BNJ08_003830, and BNJ08_005124) in the CBS10913T (Wild type) strain as described in the methods section. Since all three transporters were highly identical in terms of their primary sequences, we designated BNJ08_004188, BNJ08_003830 and BNJ08_005124 genes as *PTR_A, PTR_B* and *PTR_C*, respectively. To assess the contribution of each gene in di-/tripeptide transport we performed a growth assay wherein, only the di-or tripeptides acted as the nitrogen source for the WT and mutants. Therefore, any reduction in the cellular turbidity points to a role of the gene in peptide uptake. We used 4 dipeptides, namely Leucine-Glycine (Leu-Gly), Histidine-Leucine (His-Leu), Tyrosine-Phenylalanine (Tyr-Phe) and Glycine-histidine (Gly-His), and 2 tripeptides, viz., Phenylalanine-Glycine-Glycine (Phe-Gly-Gly) and Glycine-Glycine-Glycine (Gly-Gly-Gly) in our uptake assays. The results are presented in **Figure 5**. With the dipeptides, it is evident from the results that barring Gly-His, deletion of *PTR_A* did not result in any significant change in the growth profile compared to the WT. With Gly-His as well, the reduction in cellular turbidity was observed to be rather modest (**Figure 5D**). Contrarily, we noticed far more significant consequences on the deletion of *PTR_B* and *PTR_C*, with a major reduction in cellular turbidity observed with the Δ*PTR_C* strain. When Phe-Gly-Gly was used as the substrate, a similar profile was obtained, wherein the reduction in turbidity was of the order Δ*PTR_C*>Δ*PTR_B*>Δ*PTR_A* (**Figure 5E**). Interestingly, there wasn’t any significant difference in the growth profile of the WT and PTR mutants with Gly-Gly-Gly as the nitrogen source, plausibly indicating a similar preference for the same by all the transporters (**Figure 5F)**.

**Figure 5:**
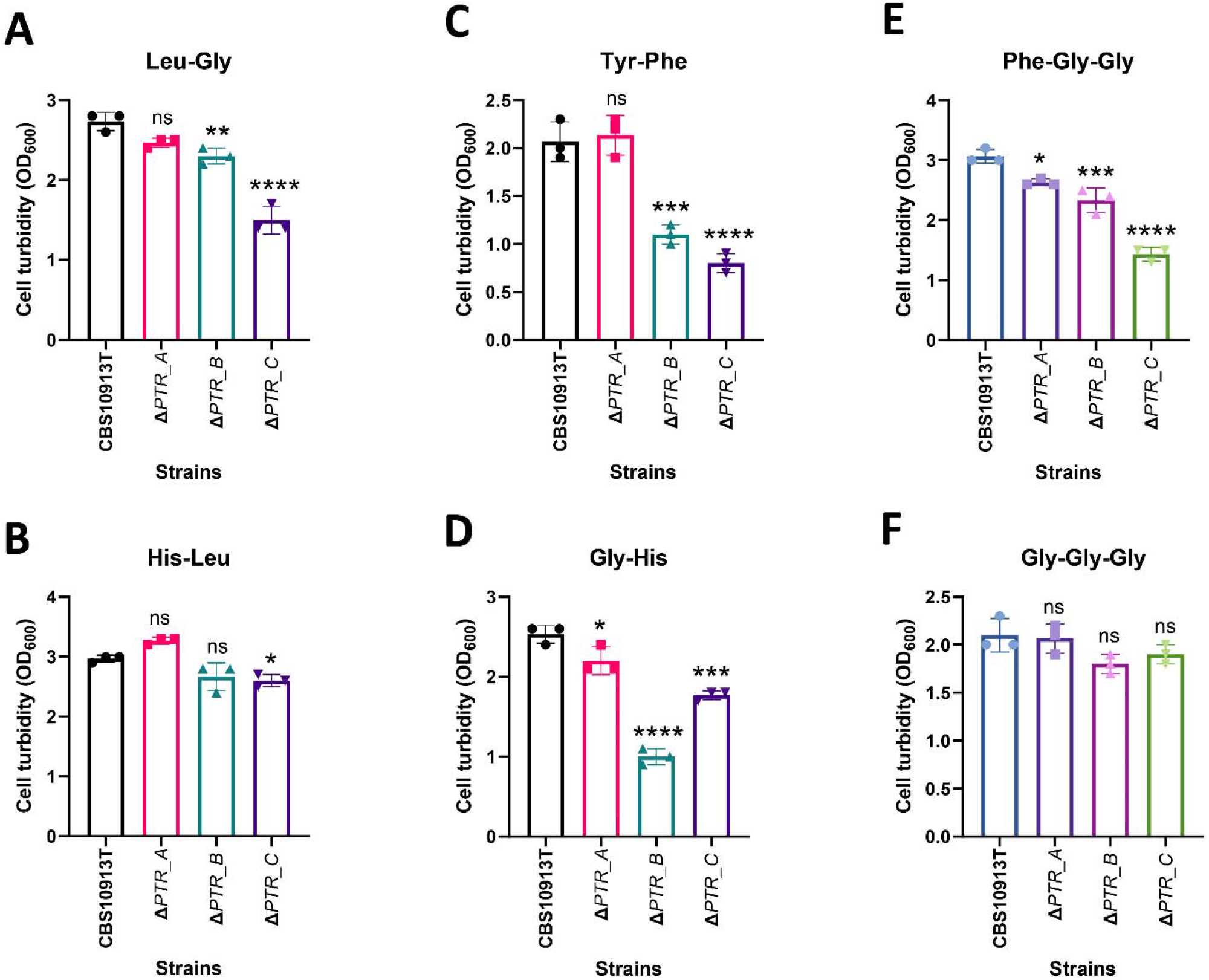
Di-/tripeptide uptake assays in the WT and PTR mutants of *C. auris*. Herein Di-/tripeptide utilization capabilities inferred through cellular turbidity served an indicator for their relative contribution in transport as elaborated in the methods. Results are expressed as Mean ± SD. One way ANOVA followed by Dunnett’s test has been applied to calculate significance. *, **, *** and **** indicate p values ≤ 0.05, 0.01, 0.001 and 0.0001 respectively.

### Assessment of Nikkomycin Z sensitivity in the PTR deletion mutants

Nikkomycin Z is a peptidyl nucleoside (**Figure 6A**) that targets the chitin synthase and has shown promising antifungal activities against many fungal species, including *C. albicans* (Larwood 2020). Of note, in a human phase I trial for treatment of coccidioidomycosis, it has shown promising results (Larwood 2020). A recent *in vitro* assessment of Nikkomycin Z efficacy against *C. auris* demonstrated an overall mode of 2mg/L as the Minimum inhibitory concentration (MIC) (Bentz et al. 2021). However, the compound was not found to be effective against all the tested isolates, the majority being the Clade III strains with MICs ranging between 16 to >64. Nonetheless, since it is quite active against the maximally drug-resistant clade I isolates, the therapeutic potential should be addressed in detail. Given the previous reports suggesting the role of PTR transporters in their uptake (Yadan et al. 1984), we chose to assess the relative contribution of the three PTR transporters encoded in the *C. auris* genome. Growth assay in liquid medium demonstrated significantly increased resistance towards Nikkomycin Z in the case of Δ*PTR_C* strain only when compared to the WT (**Figure 6B**). Notably, serial dilution spot assays on solid media also corroborated the findings from liquid media, wherein, the Δ*PTR_C* strain showed enhanced resistance compared to the WT and other PTR mutants (**Figure 6C**). However, it is pertinent to mention that all the strains showed much better growth in solid compared to the liquid medium.

**Figure 6:**
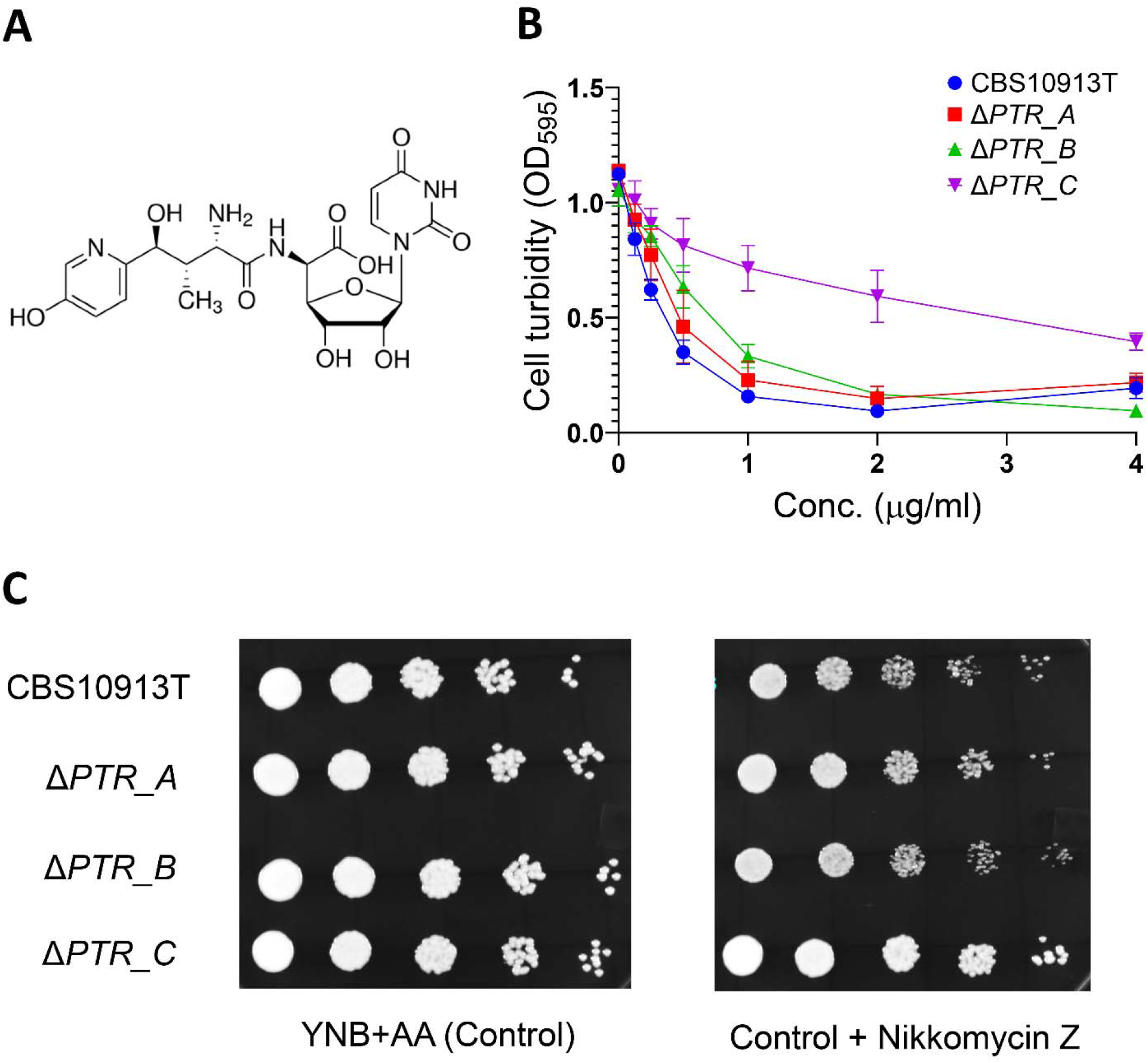
Nikkomycin Z sensitivity assays. **(A)** Chemical structure of Nikkomycin Z **(B)** Dose dependent inhibitory curves of Nikkomycin Z against CBS10913T and the PTR mutants. **(C)** Serial dilution spot assay of CBS10913T and the PTR mutants on Solid Yeast nitrogen base + amino acids (YNB + AA) medium with (*left panel*) or without 2 µg/ml Nikkomycin Z (*right panel*).

## DISCUSSION

In this study, we have identified all the putative PTR transporters encoded in the genome of five prominent *Candida* species, including the recently emerged *Candida auris*. With a significant increase in the burden of fungal infections due to *Candida* species, effective alternatives to conventional antifungal therapy are urgently needed. With demonstrated efficacy of peptide transporters in transporting peptide drug conjugates, these systems can be exploited as antifungal delivery systems. However, that would require a complete understanding of their structure and function. The present study provides a template for all such future investigations because it reports all the essential primary sequence and structure-related information for the proteins. It is clear from the study that within the PTR/POT family of *Candida* species, there are two distinct classes of transporters, where only one is significantly like ScPtr2 in terms of sequence and membrane topology. Moreover, there are few *Candida* species, namely *C. glabrata* and *C. auris* which harbor only proteins similar to ScPtr2. Given that, both CaPtr2 and CaPtr22 have earlier been shown to be different in terms of their substrate specificity, the reason and implications for such differences need to be understood for future exploitation of these proteins as drug delivery systems.

With the recent emergence of the multidrug-resistant *C. auris*, alternative therapeutics are in much demand. Under such a scenario, promising activities of peptide-based antifungal compounds like Nva-FMDP and Nikkomycin Z are quite encouraging (Bentz et al. 2021; Shahi et al. 2022). Since previous studies pointed towards a PTR transporter dependent uptake for these compounds (Yadan et al. 1984; Schielmann et al. 2017), and there are three highly identical transporters encoded in *C. auris*, we investigated the relative contribution of each transporter in di/tripeptide as well as Nikkomycin Z uptake. We constructed deletion mutants for all three genes and subjected them to di/tripeptide uptake assay and Nikkomycin Z sensitivity assays along with the WT. Overall, it could be said that compared to the other two PTR transporters in *C. auris, PTR_C* (BNJ08_005124) seems to be the more dominant player responsible for uptake of di-/tripeptides and peptide-based therapeutics like Nikkomycin Z. However, the role played by the other two genes *PTR_A* (BNJ08_004188) and *PTR_B* (BNJ08_003830) could not be ignored, more so for *PTR_B* which also showed significant differences in the di-/tripeptide uptake assays when compared to the WT. Since these proteins are highly similar to each other in terms of their primary sequence, the differential profiles observed need to be addressed in further studies through unbiased systems lacking native PTR transporters. Altogether, the study serves as a useful reference for guiding future structure-function studies on the highly relevant PTR/POT membrane transporter family.

## ACKNOWLEDGEMENTS

RK acknowledges funding support from SERB in the form of a Junior Research Fellowship under grant no. SRG/2019/000514. AB, AML, and RP acknowledge funding support from the Department of Biotechnology, Government of India through grant no. BT/PR32349/MED/29/1456/2019. SS acknowledges the Senior Research Fellowship award from the Indian Council of Medical Research, Government of India.

## DATA AVAILABILITY STATEMENT

Data available on request from the authors

