## Supplementary figure 1 and Supplementary tables 1 and 2 for "Genome-wide analysis of PTR/POT transporters in *Candida* species and their functional characterization in the newly emerged pathogen *Candida auris*"

**SUPPLEMENTARY FIGURES**

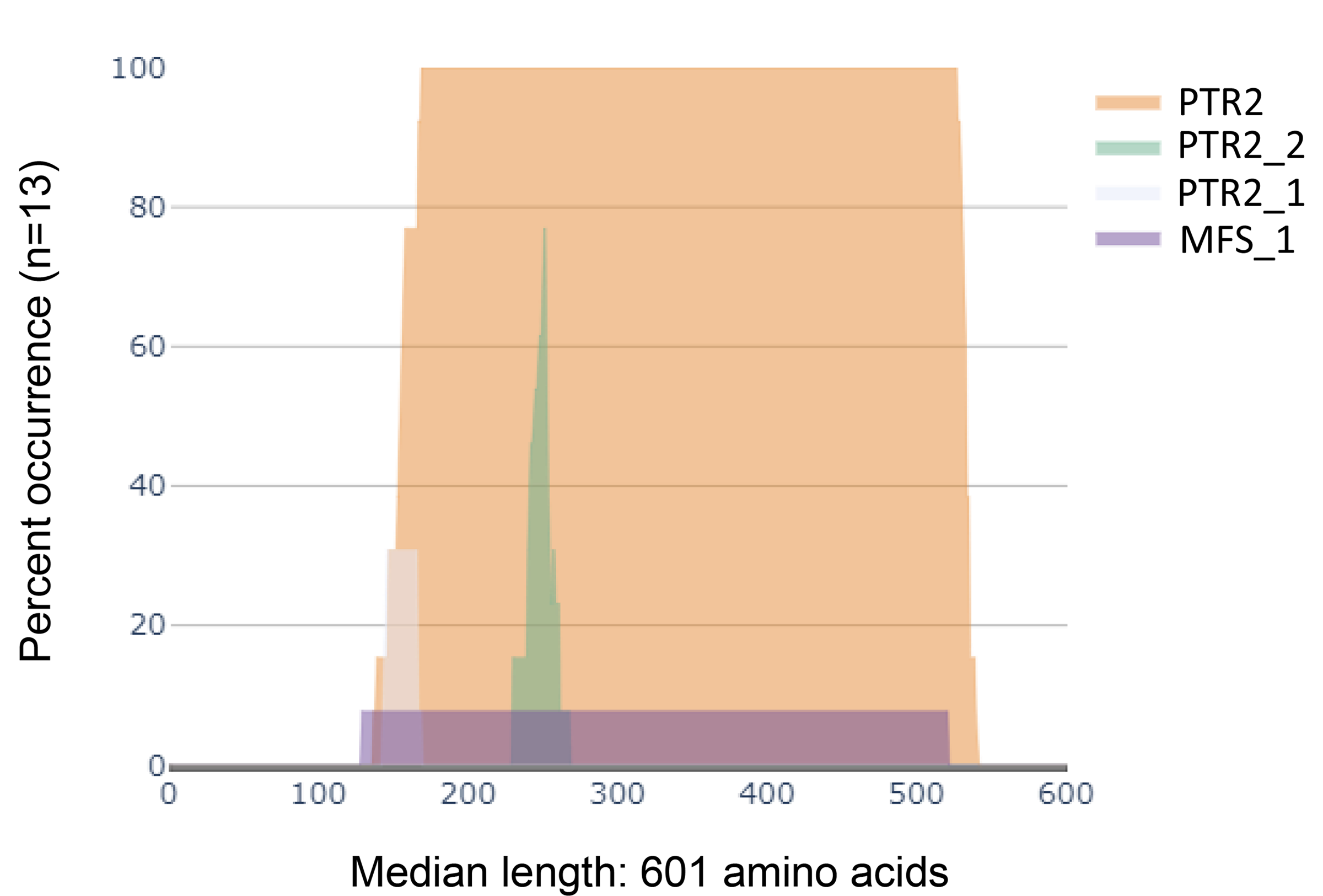

**Figure 1:** Distribution of protein domains in the identified PTR protein sequences as predicted using DomainViz. While MFS_1 and PTR symbolize the PFAM domains, PTR2_1 and PTR_2 are the PROSITE signatures.

**Supplementary table 1: List of strains used in the study**

| **S. No.** | **Strain** | **Description** | **Reference** |
| --- | --- | --- | --- |
| 1. | CBS10913T | Clinical isolate | (Wasi et al., 2019) |
| 2. | Δ*PTR_A* | BNJ08_004188 gene deleted from CBS10913T and replaced with *NAT* | This study |
| 3. | Δ*PTR_B* | BNJ08_003830 gene deleted from CBS10913T and replaced with *NAT* | This study |
| 4. | Δ*PTR_C* | BNJ08_005124 gene deleted from CBS10913T and replaced with *NAT* | This study |

**Supplementary table 2: List of oligonucleotides used in the study**

| **S. No.** | **Primer ID** | **Sequence (5'-3')** |
| --- | --- | --- |
| **Knockout construction primers** | | |
| 1. | 4188/P1 | GAGTTCTTCCACAGCAGCTGATC |
| 2. | 4188/P4 | CATCTGGAGGAATGTAGCCGTG |
| 3. | 4188/P5 | CGCTCTGTCCAACCTATCCACG |
| 4. | 4188/P6 | GCGTCGACCTGCAGCGTACGATGCTTGAGAGAAAAGAGTTGGC |
| 5. | 4188/P7 | CGACGGTGTCGGTCTCGTAGGATTACTGAGTCGTTGGCTGCTC |
| 6. | 4188/P8 | CTCAAGGTGAGTTAGACCAGCAG |
| 7. | 4188/P13 | CACTTGAAGAACTCGGGCGCCAC |
| 8. | 4188/P14 | GGTAGATAAAGTGCTGCAAGACAC |
| 9. | 3830/P1 | GGCGAAATTCGCACAAGGAAAAAC |
| 10. | 3830/P4 | GGACAACCGCTAAATCAGGTCTG |
| 11. | 3830/P5 | CGGCGCCAGCGAAAAGAAATGGC |
| 12. | 3830/P6 | GCGTCGACCTGCAGCGTACGATGACGCTGTAGTGATATTAAAT |
| 13. | 3830/P7 | CGACGGTGTCGGTCTCGTAGCACTCGTTCTTTCAAGCAGTATG |
| 14 | 3830/P8 | TGTCAAAAATGCGAATTTGCAGCC |
| 15. | 3830/P13 | GGTGAGAGGGTAATTGAGGACCC |
| 16. | 3830/P14 | CAAGGAGTCTGAAGGGCAATGTG |
| 17. | 5124/P1 | GAAAGATTGCCGTTGCCGACGATG |
| 18. | 5124/P4 | GATAGCAAGGAAGCTATGGCTCAC |
| 19. | 5124/P5 | GTCGTGTGCTACCTACCGAGGACT |
| 20. | 5124/P6 | GCGTCGACCTGCAGCGTACGATGCTAGACGTGGAAATACGAATAC |
| 21. | 5124/P7 | CGACGGTGTCGGTCTCGTAGGACGAGATCCTTCAAGGACATGA |
| 22. | 5124/P8 | GATAGAACAGTTCAGCTTTCCATC |
| 23. | 5124/P13 | GTCGCTCCCATCCATGTCTCAAAAC |
| 24. | 5124/P14 | GGTTTGGGATATTTTTGAAGCCGGG |
| 25. | NAT_P2 | TGCGCACGTCAAGACTGTCAAGG |
| 26. | NAT_P3 | TGTGAATGCTGGTCGCTATACTGC |
| 27. | NAT_P9 | CGTACGCTGCAGGTCGACgccttccgctgctaggcgcgccgtg |
| 28. | NAT_P10 | GTCTACTACTTTGGATGATAC |
| 29. | NAT_P11 | TCTGTTCCAACCAGAATAAG |
| 30. | NAT_P12 | ctacgagaccgacaccgtcgggccgctgacGAAGT |
| **Sequencing primers for knockout confirmation** | | |
| 1. | 4188_SEQ1 | GATAGTGCTGCCCCAAGCAAGACC |
| 2. | 4188_SEQ2 | GAGATCACCCTACATTTGCCACAGTG |
| 3. | 3830_SEQ1 | CTCTCCGTTACCTTTGGCGCTGTAC |
| 4. | 3830_SEQ2 | CGCTGCTAATATCTCATTACACTC |
| 5. | 5124_SEQ1 | CGCCCTGTTTAGTTGCATATGCG |
| 6. | 5124_SEQ2 | GTGAAAGCATCTTTGAACCCGACTG |
